# Plasmids, Viruses, And Other Circular Elements In Rat Gut

**DOI:** 10.1101/143420

**Authors:** Tue Sparholt Jørgensen, Martin Asser Hansen, Zhuofei Xu, Michael A. Tabak, Søren J Sørensen, Lars Hestbjerg Hansen

## Abstract

Circular DNA such as plasmids and some viruses is the major source of genetic variation in bacteria and thus has the same important evolutionary function as sexual reproduction in eukaryotic species: It allows dissemination of advantageous traits through bacterial populations. Here, we present the largest collection of novel complete extrachromosomal genetic elements to date, and compare the diversity, distribution, and content of circular sequences from 12 rat cecum samples from the pristine Falkland Islands and Danish hospital sewers, two environments with contrasting anthropogenic impact. Using a validated pipeline, we find 1,869 complete, circular, non-redundant sequences, of which only 114 are previously described. While sequences of similar size from the two environments share general features, the size distribution of the elements between environments differs significantly, with hospital sewer samples hosting larger circular elements than Falkland Island samples, a possible consequence of the massive anthropogenic influence in the hospital sewer environment. Several antibiotic resistance genes have been identified with a notably larger diversity in hospital sewer samples than in Falkland Islands samples in concordance with expectations. Our findings suggest that even though sequences of similar length carry similar traits, the mobilome of rat gut bacteria are affected by human activities in that sewer rats have larger elements and more diverse large elements than pristine island rats. More than 1000 novel and not classified small sequences was identified and hint the existence of a biological unit not previously described on a community level.

**List of figures:**

1. Sampling sites and Rat gut anatomy
2. Rarefaction curves of circular elements
3. Elements shared between environments
4. Size distribution plots of rep, mob, stab, capsid carrying elements.
5. Graphical representation of representative elements
6. Replication genes content, diversity
7. Rep_2 plasmid replication gene phylogeny
8. Table Resfams findings

**List of appendices:**

1. basic statistics table
2. Phylogeny of plasmid replication genes
3. List of known plasmids and circular sequences found in 1869 elements
4. Automatic identification of identical circular elements with different breaking points
5. Test of separation of plasmids and viruses based on predicted, annotated genes
6. No false positive circular sequences from MG1655 genomic sequencing.

## Introduction

The field of plasmid research has been revolutionized by technological quantum leaps that have allowed scientists to apply knowledge from classical microbiological studies to environmental samples. This has revealed an unseen and largely unexplained diversity of predominantly small circular elements(Lopez-Bueno *et al.*, 2009; Kav *et al.*, 2012; Minot *et al.*, 2013; Jørgensen *et al.*, 2014; Sachsenröder *et al.*, 2014). It is imperative to study plasmids, viruses and other circular extrachromosomal elements as they are the main vectors in the evolution of bacteria and important in the efficient treatment of human infections with antibiotics(Barkay and Smets, 2005; Sørensen *et al.*, 2005; Gillings, 2013; Martínez, Coque and Baquero, 2014). Yet very little is known about these elements in the vast majority of microbial species that has never been cultured. Previously, circular DNA in natural environments was studied using indirect methods such as typing(Carattoli, 2011), targeting known sequences (Carattoli *et al.*, 2005; Brolund *et al.*, 2013) and capturing approaches(Jones and Marchesi, 2006; Klümper *et al.*, 2014), but with recent advances, it is possible to gain insight on the most basic level of information: The unabridged sequence of significant numbers of plasmids, viruses, and any other circular genetic element residing in a sample(Lopez-Bueno *et al.*, 2009; Li *et al.*, 2012; Walker, 2012; Kav, Benhar and Mizrahi, 2013; Sentchilo *et al.*, 2013; Jørgensen *et al.*, 2014; Sachsenröder *et al.*, 2014; Kraberger *et al.*, 2015). Though manual curating is still a fundamental step in some new plasmid recognition pipelines(Lanza *et al.*, 2014), the sheer volume of future datasets consisting of thousands of sequences, billions of reads and trillions of nucleotides will likely demand fully automated analysis pipelines.

The term plasmid is an umbrella for many unrelated genetic structures that share the feature of being cytoplasmic replicating extrachromosomal DNA molecules of secondary importance for bacterial hosts relative to chromosomal DNA(Summers, 1996). The largest known plasmids are called megaplasmids and consist of millions of nucleotides that carry hundreds of genes and can be considered as an instrumental part of the genome of the host organism(Barnett *et al.*, 2001; Finan *et al.*, 2001). Contrary, the smallest known plasmids are below one kilobase (kb) long and encode a bare minimum of genes, sometimes only a single gene needed for replication(Yasukawa *et al.*, 1991; Burian, Stuchl-¦¦ük and Kay, 1999). The term cryptic plasmid is often used to describe that a host beneficial role is not obvious as for stereotypical plasmids (Novick *et al.*, 1976). Cryptic plasmids has hitherto been a curiosum known to exist but never explored as the type have little to offer clinical or biotechnological plasmid research on antibiotic resistance plasmids. Potentially, this class could make up a significant fraction of the so-called bacterial dark matter(Rinke *et al.*, 2013) and are in some cases regarded as molecular parasites, resembling the popular idea of the selfish gene(Dawkins, 2006). Although little research has focused on these trait-less elements, entire classes could exist and be important for bacterial ecology and evolution, for example resembling satellite viruses or viroids, the smallest stable genetic entities known(Krupovic and Cvirkaite-Krupovic, 2012; Jørgensen *et al.*, 2015; Kraberger *et al.*, 2015) or by mediating horizontal gene transfer. As a parallel to the term viroid, we here suggest the term plasmoid to describe circular extrachromosomal DNA that does not fit the categories plasmid, virus, or viroid.

Despite technological advances in mobilomics, major issues remain. These issues involve the ability to physically separate mobile DNA from chromosomal, correct assembly of troublesome repetitive sequences and capacity of identifying unbiased abundance relations between plasmids and the chromosomal background. In particular, the ability to assemble sequences scarred by recombination events is problematic for dynamic structures like plasmids and viruses, making many interesting sequences go undetected by current methods. The difficulties of obtaining a bias free sample of extrachromosomal DNA are many and reviewed more thoroughly in (Jørgensen *et al.*, 2015).

The brown rat (*Rattus norvegicus*) has spread with merchant ships across the globe and is now considered a problematic invasive species responsible for decline and extinction of other species worldwide, which has been documented in the pristine Falkland Islands (Fig.1A) (Robinson, 1965; Hilton and Cuthbert, 2010). Furthermore, large populations of brown rats exist in the sewers of urban areas in temperate climate, living in close contact with human wastes (Fig.1C). Rats are able to transmit a wide range of zoonoses, why ongoing efforts seek to eliminate them from the daily lives of the human population(Meerburg, Singleton and Kijlstra, 2009; Himsworth *et al.*, 2013; Tabak *et al.*, 2014, 2015) [33,34]. Brown rats are hindgut fermenters, using a diverse microbial flora in their enlarged, pouch-like cecum to degrade fibrous materials such as woods(Yang, Manoharan and Young, 1969; Montgomery and Macy, 1982) (Fig.1B). Depending on available food sources, the brown rat will ingest any organic material, such as bird eggs and small shrubberies that dominate vegetation on the Falkland Islands or human food waste carried by sewage in urban environments. Since hospital sewers are considered a hotspot for horizontal gene transfer and rats are important zoonosis vectors(Jacobsen *et al.*, 2007; Meerburg, Singleton and Kijlstra, 2009; Czekalski *et al.*, 2012), we expect hospital sewer rats to be an optimal environment for transfer events in a gastrointestinal setting similar to that of humans, from which significant knowledge on enteric horizontal gene transfer can be derived. We hypothesize that the environment which a rat inhabits, is shaping the landscape of MGEs in the gut of the animals.

**Fig. 1.**
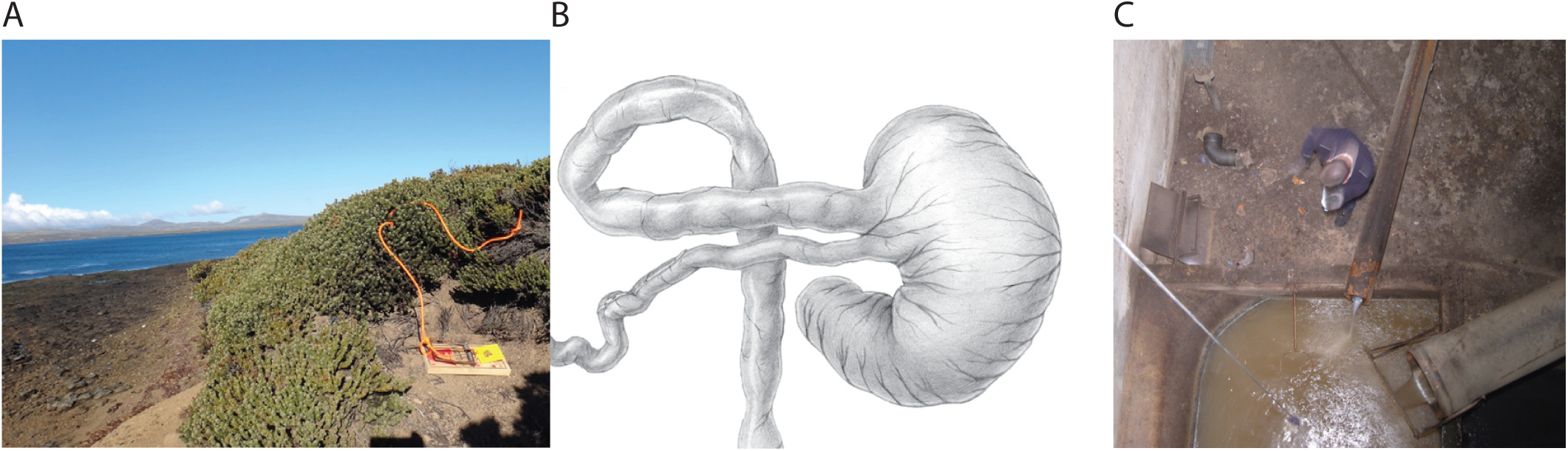
**A**, Sampling site at the Falkland Islands. The vegetation is dominated by small shrubberies that along with bird eggs and carcasses is thought to be important food sources for the Falkland Islands rats. Fig.1 B, schematic overview of the rat cecum. In the lower left, the ileum leads to the cecum, separated by a sphincter. The big, pouch-like cecum is pivotal in degradation of fibrous food items such as wood. The cecal content passes through the colon where nutrition in the shape of short chain fatty acids, an important product of cecal fiber degradation, is utilized by the rat. Fig.1 C sampling site at a sewage collecting room directly below a large hospital in Copenhagen, Denmark. Fibrous material is thought to be very limited whereas human food and waste product are thought to make up the diet of the inhabiting rat (middle left).

In this work, we describe the mobilomes of 12 samples in the largest such study to date using a validated pipeline to identify the unabridged sequence of thousands of novel plasmids, viruses and plasmoids. We demonstrate that the type of elements found in this study is far more diverse than hitherto recognized, and we provide a large database for researchers of different fields to recognize circular elements that would otherwise go unnoticed. We finally proceed to identify significant differences in the landscape of circular elements between samples from environments of contrasting anthropogenic activity.

## Results and discussion

### Sequencing statistics and chromosomal contamination

From the equivalent of three lanes on an Illumina HiSeq2000 sequencer, reads were generated, downloaded and quality trimmed yielding 618,750,653 reads with a mean quality score of 35.8 and a mean length of 105 nt. A detailed view of samples and reads can be found in appendix 1. Reads were assembled into contigs for each sample individually and on average 648 contigs larger than 1kb was found in a sample. Circular elements were identified and constitute an average of 33% of all contigs larger than 1kb, but up to 62% in a single sample (Appendix 1). This further confirms that the circularization process is efficient in finding a large part of the circular elements in a mobilome dataset, particularly as unclosed plasmids with repetitive sequences would tend to break into multiple pieces inflating the number of seemingly linear contigs. By mapping reads back to circular sequences, estimates of contribution to assembly could be estimated and is in all cases less than 10%, meaning that the large majority of sequencing reads do not contribute to assembly of contigs longer than 1kb (data not shown). This is in concordance with findings from mobilome sequencing on the PacBio RS platform, where only very few error corrected reads was not short-repeat sequence, thought to arise from amplification noise or sequencing error rather than being biological entities though this possibility cannot be excluded entirely(Jørgensen *et al.*, 2015). Data from a recent publication indicate that <10% of mobilome reads can be assigned to known plasmid and chromosomal sequences, which is consistent with the above finding, although not directly comparable (Norman *et al.*, 2014).

In order to estimate the level of chromosomal contamination, we used the hidden Markov model based rRNA_hmm script(Li, 2009) to map a subset of reads and normalized the result to number of input reads. By assuming an average copy number and genome size of an *Escherichia coli* with 4 copies and a 5M nt genome, we approximated the chromosomal contamination in the 12 samples used in this study. Six of the samples have a chromosomal contamination of 2% or lower and the other six samples had between 5 and 20 % chromosomal DNA (Appendix 1). A potential reason for the relatively high fraction of residual chromosomal DNA after prolonged exonuclease treatment could be existence of un-nicked genomes in the samples. Potentially, this could be resolved by adding a rare-cutting restriction endonuclease step to the protocol. Further, mitochondrial DNA from the rat host known to exist in the DNA purifications will inflate the calculated fraction of chromosomal DNA as rat mtDNA carry 16S rDNA. The level of chromosomal DNA is not thought to interfere with analysis of circular elements apart from lowering the total number of mobilome derived reads as we expect chromosomally derived longer contigs would be rare due to low coverage. Further, analysis of E. coli DNA show that assembly of genomic DNA in a known genetic background do not yield any false circular contigs that are actually long direct repeats or other problematic structures, indicating that chromosomal contamination is not problematic when limiting investigations to circular contigs (Appendix 6)(Jørgensen *et al.*, 2015).

### Basic circular element statistics

From 12 samples, a total of 2,272 circular elements were identified. Of these, 1,869 unique circular elements were left after replicate identification (see below, Appendix 4). Previously, we have used inverse PCR to demonstrate that at least 95% of bioinformatically identified circular elements have circular topology, substantially adding to the claims made in this study. The size range of non-redundant circular elements is 899nt to 33,547nt with an N50 value of 2862. The histogram in Fig. 4A reveals a significantly different size distribution between environments (t-test, P<0.001) and a much higher diversity of small sequences than of large sequences with 1188 elements shorter than 2kb and 681 elements longer than 2kb, of which, only 59 elements are longer than 7.5kb. Notably, while elements found in Falkland Islands (FK) samples are by far most prevalent up to 2.5kb, hospital sewer elements are more diverse in all larger size bins, despite a smaller total sequence pool (1093 and 810, respectively) and fewer biological samples (7 and 5 respectively). Here, it should be emphasized that abundance relations are obscured by the MDA amplification, meaning that we can only make qualitative and not quantitative inference on the genetic elements. Of the predicted genes annotated using the PFAM database, 2,120 ORFs out of 5,344 could be assigned to PFAM families.

Because any circular element regardless of origin could be identified in a mobilome sample, we mapped all circular elements on the complete rat genome, Rnor6.0, as rat DNA could be a likely source of eukaryotic circular DNA, e.g. from certain cancers where circular DNA is known to be formed(Storlazzi *et al.*, 2010). Complete rat mitochondrial DNA and nine full length sequences with only very few base substitutions were found in non-repetitive regions in samples from both the Falkland Islands and from hospital sewer. The rat derived circular sequences have lengths varying between 932nt and 1825nt, mostly with breakpoints different from the organization in the rat genome. While the biology of these rat derived circular elements is interesting, they will not be discussed in greater detail in this study. Notably, very few circular elements had partial hits on the rat chromosome, suggesting that rat individual genetic variation does not play a major role in this effort to exclude rat derived sequences.

### More than 100 database plasmids identified in mobilomes

We have observed that a plasmid existing in multiple mobilome samples will linearize in a different, seemingly arbitrary place every time, possibly in a weakest link fashion in a random place of lowest coverage. If circular elements with different breakpoints were compared using e.g. BLAST, the best hit would not be of total length even though the sequences are in fact identical. For this reason, it is important to take into account circularity when comparing mobilome data. Using a BLAST-based (Altschul *et al.*, 1997) search script, we identified complete, known plasmids in the dataset of 1869 circular elements (see Appendix 4 for scripts and descriptions). Within these, a total of 114 elements were close to identical to database versions (summarized BLAST hit e-value <1E-4, subject-query size difference max. 100nt). Of these 114 known elements, 90 are from a previous rat gut mobilome publication that was based on a rat sample from a different sampling site in Copenhagen prepared in another laboratory almost two years earlier, making contamination highly unlikely (Jørgensen *et al.*, 2014). Re-finding 90 elements from the very deeply sequenced (160M PE reads) mobilome in the current dataset strengthens the hypothesis that the identified circular elements are widespread in the gut of rats across time and space and raises the question if the elements could also be found in other habitats than rat gut. In addition to the 90 mobilome plasmids and circular elements, some 20 known plasmids were also identified along with *Muridae* mitochondrial DNA and three sequences linked to prion disease, found in bovine milk, bovine serum and multiple sclerosis-Affected Human Brain Tissue [43,44]. A list of known plasmids can be seen in Appendix 3. The plasmids on the list range from 1309 nt to 6935 nt and typically come from easily cultured, well described bacterial genera such as *Escherichia*, *Shigella*, *Bifidobacterium*, or *Salmonella*, and we consider this tendency to identify elements from well-studied microorganisms an illustration of the expected database bias. The fraction of known elements found in RH samples are significantly higher than FK plasmids (T-test, P=0.001, data now shown), further suggesting database bias, not just towards potential human pathogens or commensalists, but also towards well studied environments.

### Sequencing depth estimation by rarefaction

To visualize the sequencing depth of individual samples, we created random subsets of all 12 rat cecum samples, assembled the subsets and identified circular elements within contigs. In Fig.2, this effort is summarized, with red lines symbolizing hospital sewer samples and green lines symbolizing Falkland Island samples. The wavy course of the graphs are thought to be caused by low number of replicates (n=1 for each rarefaction point). The reason for this is the calculating power going into each rarefaction point. For all except one sample, a plateau of the number of circular elements and of cumulative circular sequence length is reached indicating that the vast majority of complete circular sequences that can be identified with this methodology have been identified (Fig.2A and B). For both measures, Hospital sewer samples cluster closer together than Falkland island samples. In a previous publication using a similar approach on a similar sample, a similar saturation of total contig length was seen at a rarefaction subset size of ca 50M reads, further establishing that the dataset sizes used in this study is sufficient as very little was gained between 50M and 160M reads (Jørgensen *et al.*, 2014).

**Fig. 2.**
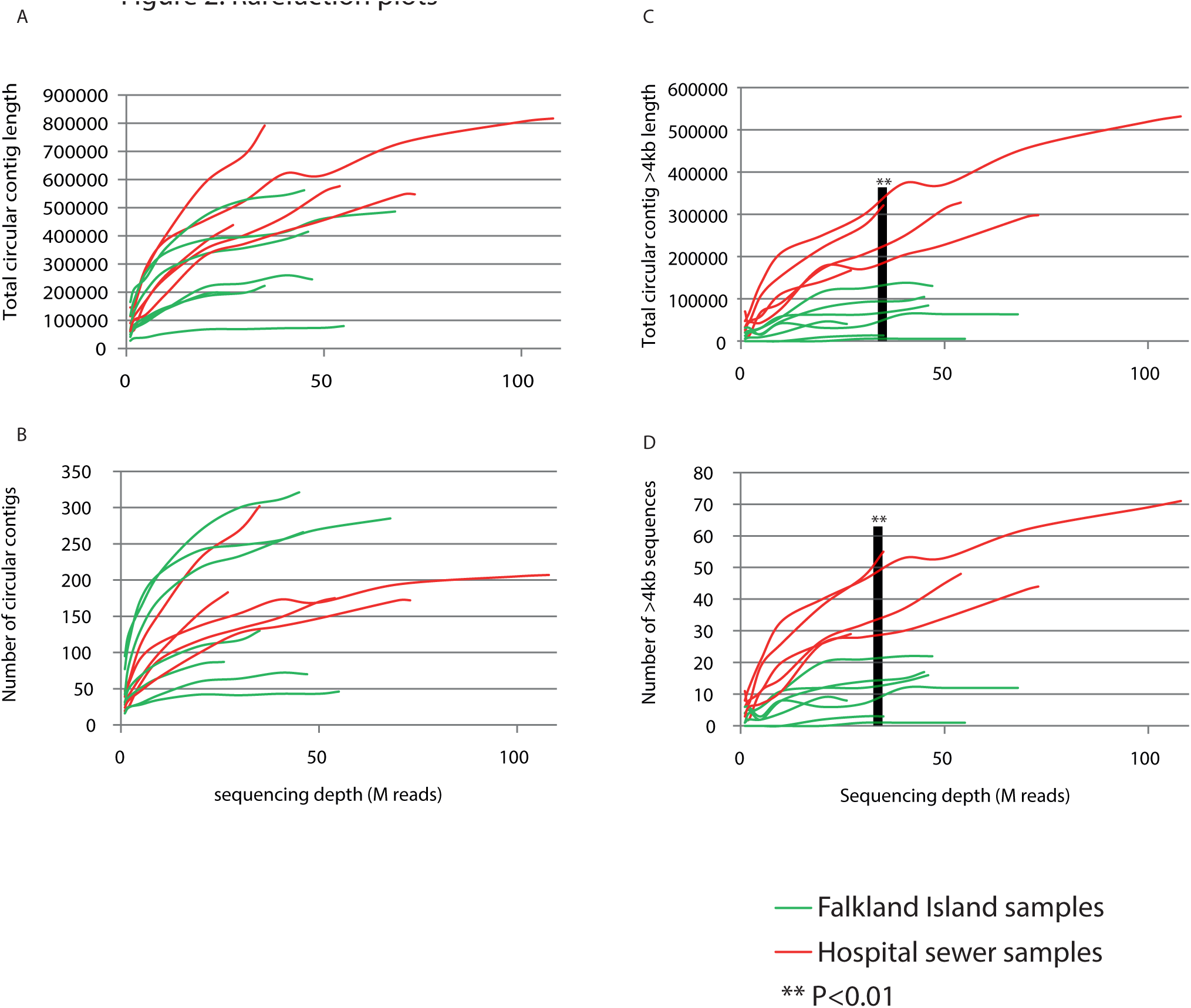
**A-D** rarefaction plots of complete, circular elements. Basic statistics of circular elements found in each sample is plotted as a function of the size of the dataset used (M reads used for assembly). A and C shows the total number of nucleotides in circular elements counting either all elements (**A**) or elements longer than 4 kb (**C**). B and D show the number of circular elements counting either all elements (**B**) or elements longer than 4kb (**D**). For neither metric, a difference between environments can be seen when observing all elements, but for elements longer than 4kb, a pattern of hospital sewer samples having more elements and a longer total contig length emerge. This observation is supported by t-test that is showing a significant difference between environments (p<0.01).

There is no significant difference in the cumulative circular sequence length or in the number of sequences between the environments when counting all contigs, but if sequences longer that 4kb are extracted, a significant difference is seen (Fig.2C and D). Here, both number of elements and cumulative sequence length are greater for hospital sewer samples than for Falkland Island samples (t-test, P<0.01 for both) with no Falkland Island samples having higher sequence diversity (Fig.2D) or cumulative sequence length (Fig.2C) than the most anemic hospital sewer sample. The 40M read subset was used for this evaluation to avoid sampling size bias from hospital sewer samples with >70M reads, as illustrated with a grey bar in Fig2C and D. The higher diversity of long sequences from hospital sewer samples than from Falkland Islands samples could indicate an anthropogenic effect. Thus, longer sequences are thought to be more likely to harbor accessory genes (e.g. resistance genes) that might be of greater importance in a heavily polluted environment like a hospital sewer than on uninhabited islands. From phylogenetic assignment of 16S rDNA amplicons from 27 samples, we know that the phylum level distribution is similar across biological replicates and environments and that an average of 25- 30% of OTUs is shared between samples (see manuscript 4, Fig.4B+C). Between certain samples, half of OTUs are shared. While we are not able to identify the hosts of the plasmids, viruses and plasmoids, we assume that the relatively stable phylum distribution and large fraction of shared OTUs across samples do not fundamentally influence the length or composition of the pool of elements that are able to exist in a sample.

### Plasmids, viruses and plasmoids are highly individual between rats

In order to quantify overlaps in MGE diversity between samples and environments, the same approach as for identifying known plasmids was used (see above and Appendix 4). The four measures of circular element redundancy seen in Fig.3 reveal a closer kinship of hospital rat mobilomes than of Falkland Island rat mobilomes. In non-redundant circular element pools of environments, 1.8% of sequences are shared between Falkland Island and hospital sewer samples (34 sequences), as visualized in the area-proportional Venn-diagram in Fig.3A. While this is a small fraction, the existence of identical sequences found in contrasting environments separated by thousands of kilometers with no human vector of spread points to a widespread pool of circular entities that currently unrecognized. Unsurprisingly, the overlap of 73 sequences between hospital sewer samples from this study and a hospital sewer sample from a previous study is larger than the overlap with Falkland Islands samples of 34 sequences, even though the two hospital sewer datasets are from different sampling sites within Copenhagen (Jørgensen *et al.*, 2014).

**Fig. 3.**
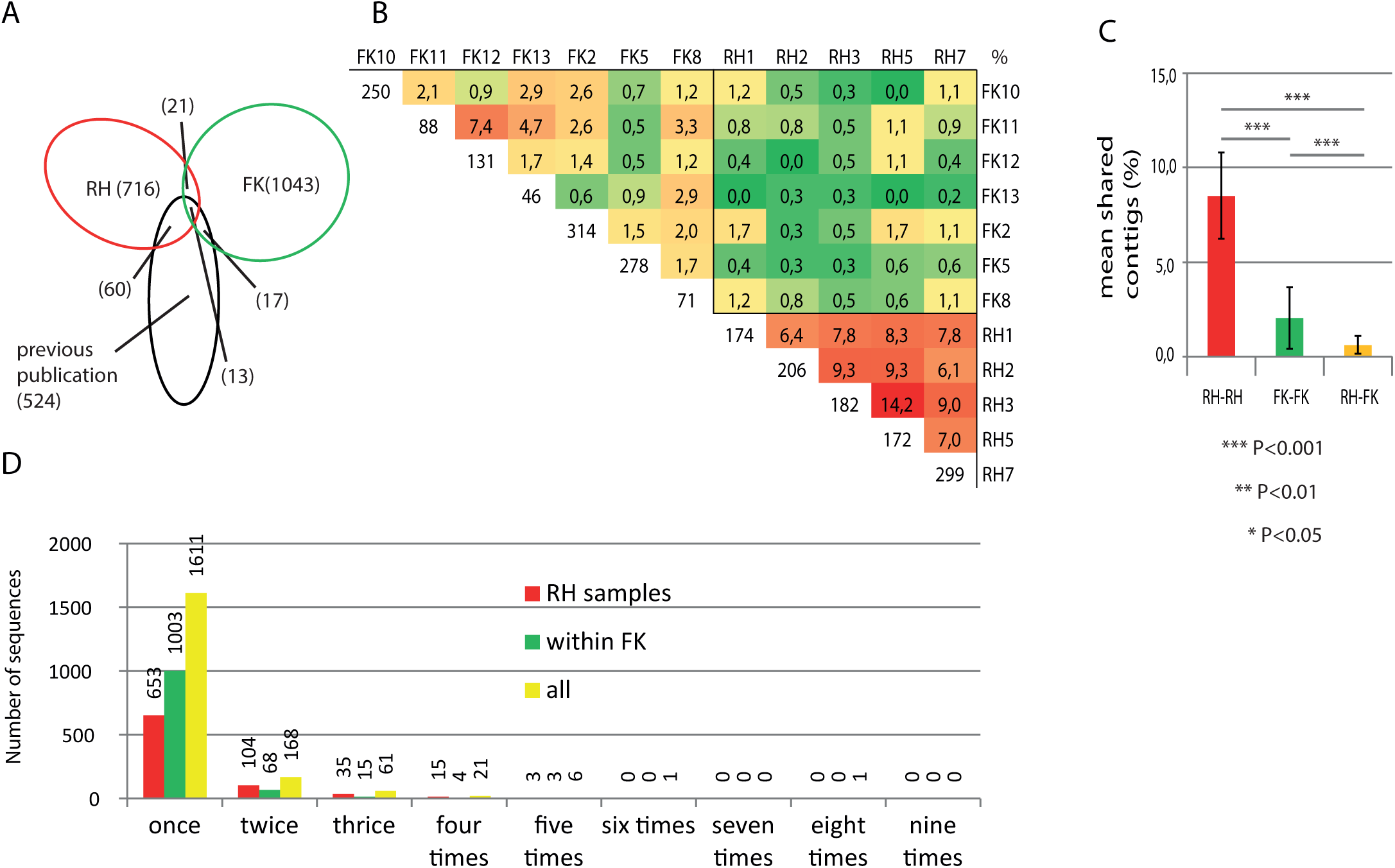
**A**, area proportional Venn diagram showing overlap between the elements identified in this work (red and green) and in previous work (black). Multiple elements are found in two or three of the environments suggesting a presence of elements across time and space. The large majority of elements, however, is only found in one environment. **B**, Jaccard similarities (percent of shared elements) for all combinations of samples with the total number of circular elements identified displayed in the bottom cell of each column. The colorization and the summation of average numbers in **C** show that hospital sewer samples (RH) are significantly more akin than Falkland Island samples (FK). Further, a significantly smaller Jaccard similarity is observed between environments than within them, suggesting that each environment has a distinct population of elements and a smaller shared population of elements. In **D**, the number of samples each non-redundant element is encountered in is summarized, with by far most sequences encountered only once. With the large number of identified sequences, still hundreds are encountered more than once, with e.g. a single element encountered in eight out of 12 samples.

The heatmap of Jaccard similarity in Fig. 3B illustrate all possible sample-sample distances, with indication of percent shared identical circular elements. The color intensity reveals a significantly higher similarity between RH samples than between FK samples, an observation that is statistically confirmed by the summation of average identical sequences between samples in Fig.3C (t-test, P<0.001). Further, a significantly smaller fraction of elements were identical between environments than within them (t-test, P<0.001). This is expected and likely caused by factors like geographical distribution and divergent habitats, but surprisingly, the fraction of elements found more than once is low even for RH specimens caught within meters from each other (258/1869, Fig. 3B and D), hinting at a largely individual diversity of circular elements in the rat cecum. In both environments, it is most common for elements to be found in only one sample, making up a total of 1611 out of 1869 elements. One element is found six times and one element is found eight times, indicating that these elements are widespread despite of the geographical separation of the Atlantic Ocean and some 13,000km.

### PFAM based plasmid and virus separation

The applied methodology allow any circular DNA present in the original samples to be amplified and characterized, including ssDNA and dsDNA of prokaryotic, eukaryotic, and viral origin (Sachsenröder *et al.*, 2014). For this reason, we expect a mixture of plasmids, viruses and plasmoids to be present in the dataset of 1869 sequences. We set out to separate them, based on complete, predicted genes annotated using HMMER3 and the PFAM27 database. To evaluate how efficient the separation of plasmids and viruses was, we tested it on a mock community of plasmid and virus sequences downloaded from NCBI. See Appendix 5 for complete procedure and validation. The test showed that separation is close to complete, with very few sequences carrying plasmid-like genes being wrongly annotated (0.25%). Ca 20% of sequences could not be annotated using this approach, possibly owing to incomplete PFAM family lists or simply sequences without classical plasmid or viral protein coding genes. This fraction is expected to be significantly higher in a mobilome than in a collection of curated, well defined plasmids and viruses. A significant number of database plasmids were found to carry capsid gene(s), accounting for 1/4^th^ of all identified capsid protein and possibly reflecting phages integrated on plasmids. This way, it is possible to separate plasmids and viruses, if taking into account integrated phages on plasmids.

By analyzing PFAM assignment of genes to plasmid and virus traits in the 1869 non-redundant circular elements, we found 504 plasmid-like elements, of which, 4 also carried a viral capsid protein. Further, 120 virus-like elements and 1,245 elements without identified plasmid or virus traits was found. The elements have all been annotated and are available at EBI under the study number PRJEB8617.

For the four subgroups plasmid replication, plasmid mobilization&stability, virus-like, and plasmoid, the size distribution is significantly different between the environments (t-test, P<0.05-0.001), indicating an anthropogenic effect on circular sequence size and composition (Fig.4B-E). For sequences carrying plasmid-like genes, the significantly different size distributions are not obvious at first sight (Fig.4B and D, P<0.01 and P<0.05, respectively). In both cases, the majority of sequences carrying the plasmid-related trait are between 2kb and 5kb. Interestingly, sequences that carry a plasmid mobilization or stability gene are in average longer than sequences that carry a plasmid replication gene. This is in accordance with expectations as replication is an essential feature in a plasmid whereas stability and mobilization are secondary features that will rarely stand alone as replication genes are able to.

**Fig. 4.**
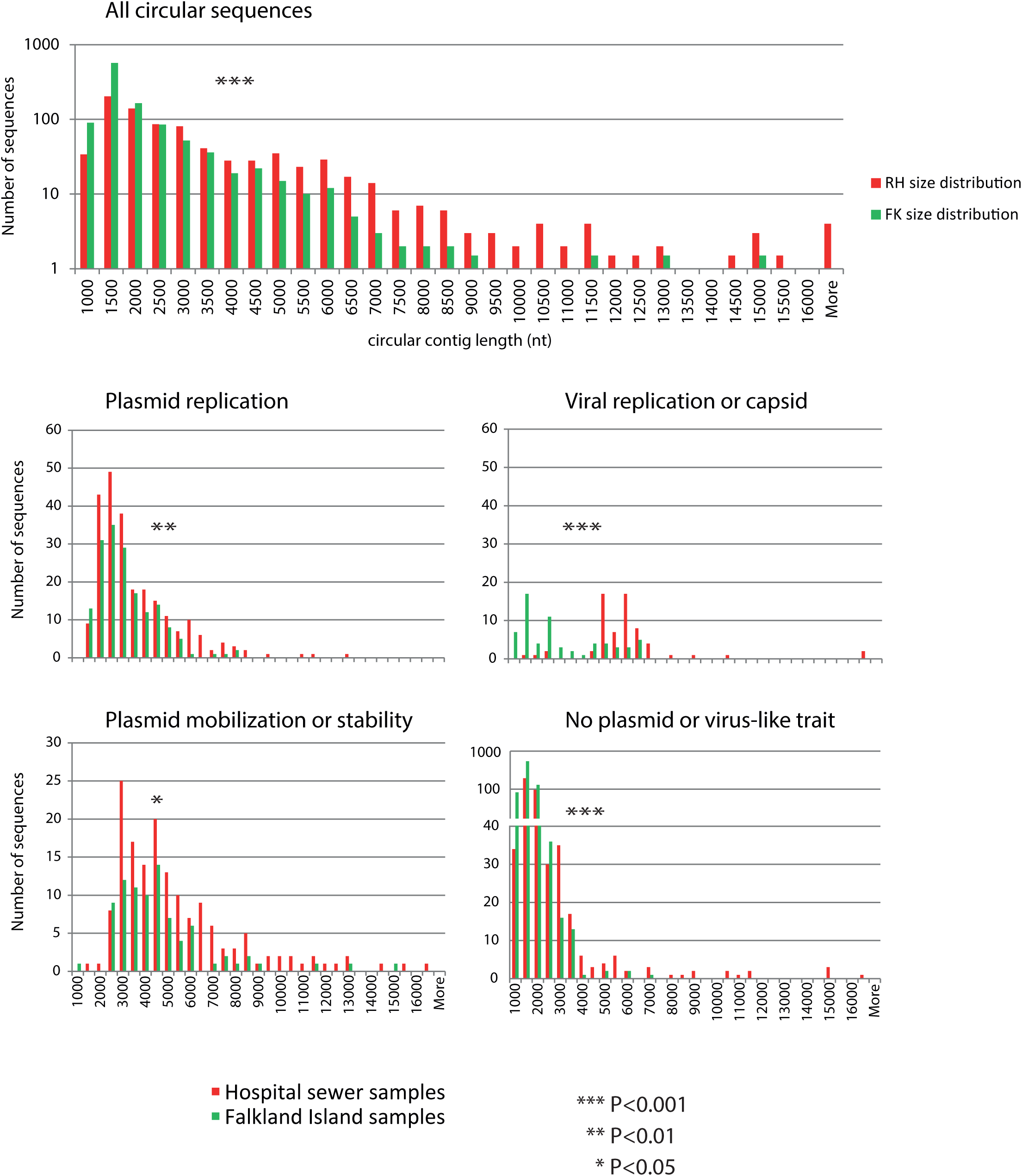
Size distribution plots of circular elements with RH denoting hospital sewer samples and FK denoting Falkland Islands samples. In **A**, a logarithmic scale is used to demonstrate the total number of nonredundant circular elements in either environment, with a significantly different size distribution. Likely, this is driven by the large number of small plasmid or virus. Notably, the vast majority of short elements <2kb (see **A**) can be found in this category, with only very few elements longer than 4kb, highlighting that very few sequences of more than 4kb cannot be categorized as either plasmid or virus. To achieve sufficient visualization of short and long sequences, the chart in D is a composite between linear and logarithmic scales, as marked on the axis.

**Fig. 5.**
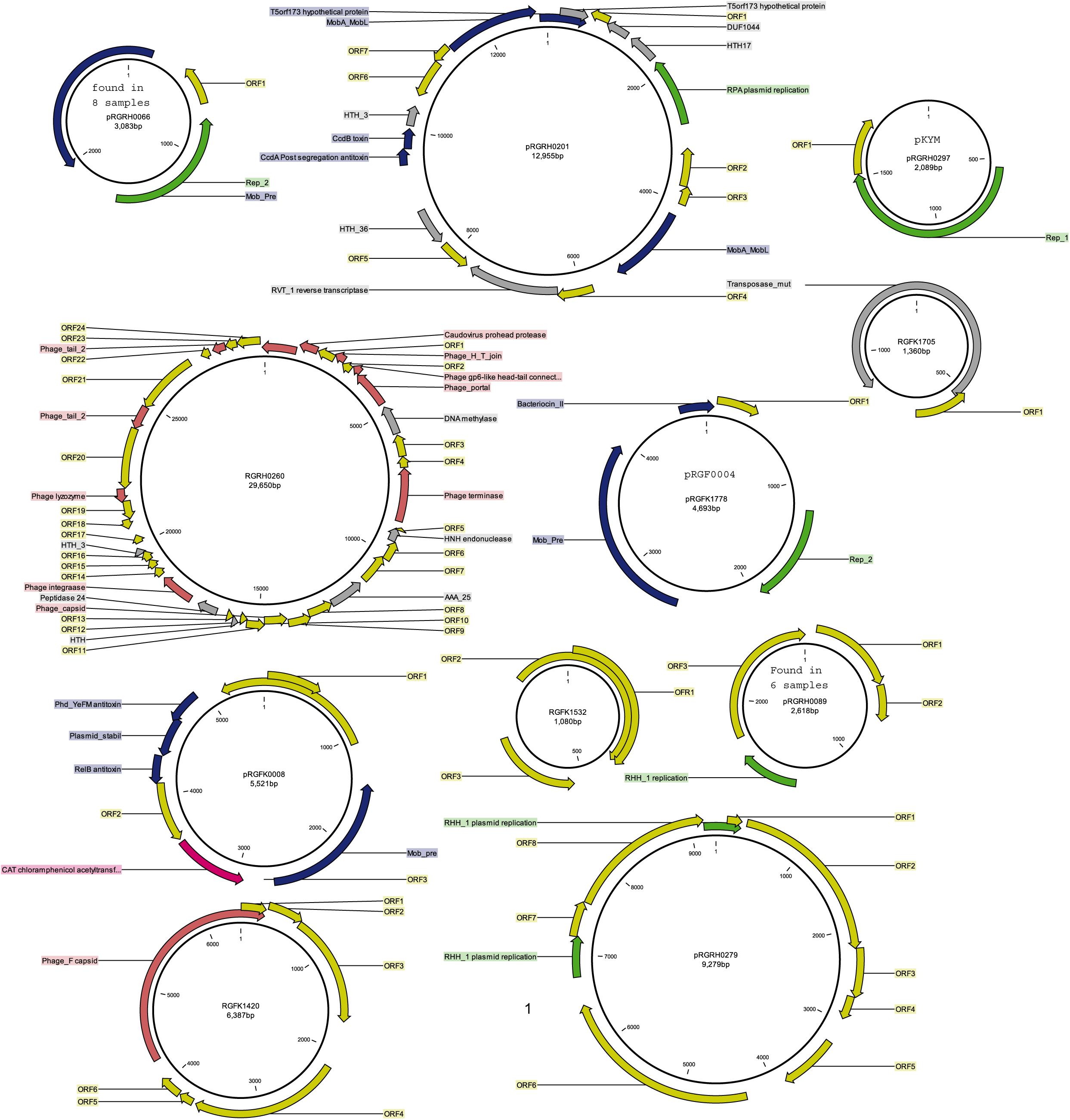
graphical representation of 0.59% of the identified elements selected to represent the types of element found: stereotypical plasmids, viruses, cryptic plasmids and plasmoids. Further, one of the depicted elements have been found in eight of 12 samples, one is a known plasmid (pKYM) and one is identical to an element identified in a previous publication (pRGF0004).

The by far most pronounced difference between environments is the size distribution of circular sequences carrying a viral replication or capsid gene (Fig.4C). Here, two distinct distributions with maxima at 2kb and 5.5kb are clearly and significantly dividing the environments. While both environments have sequences in both groups, the majority of FK sequences are within the 2kb distribution whereas the majority of RH sequences are within the 5.5kb distribution leading to speculation on environment selection, but closer inspection reveal that the 2kb maximum is almost exclusively derived from a single sample from the Falkland Islands and thus possibly attributable to individual variation rather than environment selection (data not shown).

The size distribution of sequences without recognizable plasmid or virus-like traits (Fig.4E) is heavily biased towards small sequences, demonstrating that few sequences above 4kb cannot be classified based on their genetic content. Notice that the scale for Fig.4E changes from linear to logarithmic at 40 sequences to allow representation of all groups, emphasizing the diversity of very small molecules.

In an attempt to decipher the differences in environment distribution of plasmid and virus-like traits, we analyzed the sequences in two size-fractions chosen to represent the sequences above and below that of the smallest known plasmids: above 2kb and below 2kb (Fig. 6B). The difference in frequencies of plasmid or virus related traits between these groups is striking: for sequences smaller than 2kb only ca 10% of sequences harbors a gene with plasmid or virus-like domain whereas this is case for ca 70% of sequences longer than 2kb. Analyzing plasmid database sequences yields a similar ca 70% assigned sequences (see Appendix 5), indicating that the applied separation is not able to identify all plasmids and viruses. It is surprising that most large sequences can be classified because the mobilomes approach is independent of database bias and it is an indication that few completely unknown classes of plasmid or virus backbone genes remain to be identified in rat gut, at least for sequences above 2kb. This finding goes against a primordial belief among many microbiologists: that a huge diversity tail of rare species/sequences exists in microbial communities. However, even though most larger sequences can be classified based on PFAM annotation, the nucleotide composition vary greatly as shown previously for replication genes in a mobilome (Jørgensen *et al.*, 2014).

**Fig. 6.**
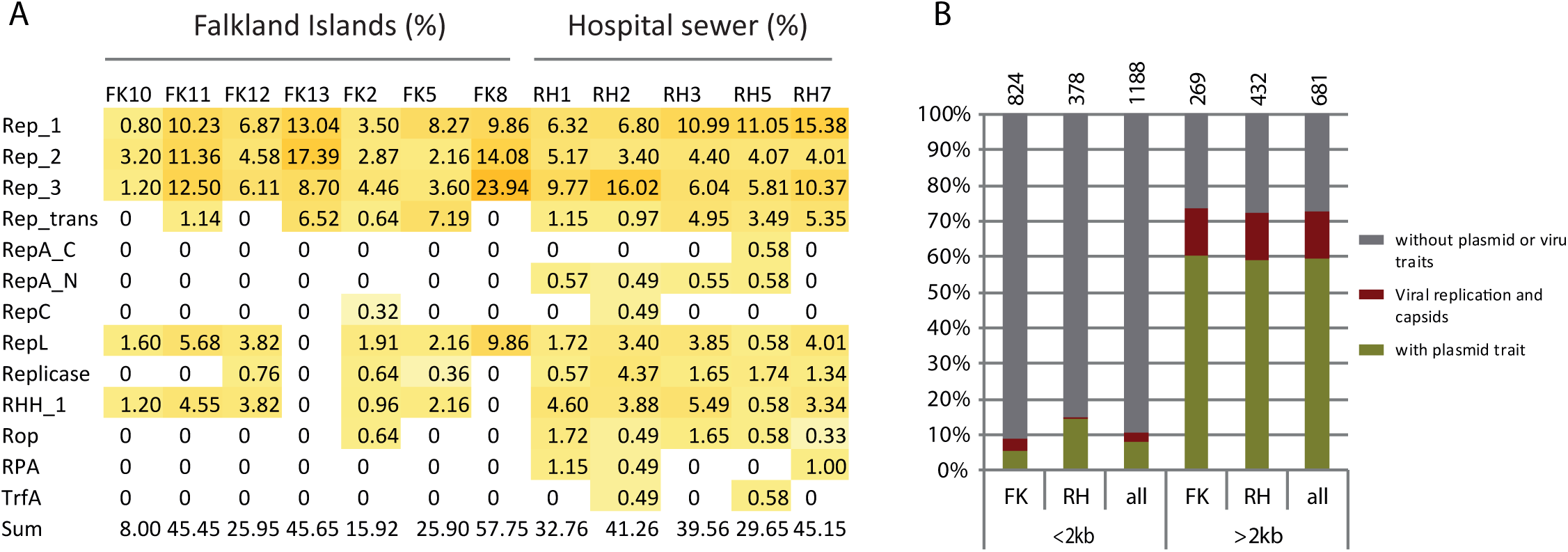
**A**, replication gene diversity and fraction. Numbers correspond to percent of elements from each sample carrying a plasmid replication gene, summed in the bottom row. Between 8% and 57% of elements carry a replication gene, mostly from the families Rep_1, Rep_2, and Rep_3. **B**, division of all elements by size reveals a fundamental difference in the fraction of sequences assignable to either virus or plasmid bins. Where only ca 10% of elements <2kb long can be assigned, this is the case for 70% of longer, >2kb elements, suggesting that

Interestingly, the difference between environments in circular sequence composition seen in fig.4B and D is completely cancelled out by the size-binning. This way, the difference between Falkland Island and hospital sewer samples lies in the size distribution of sequences rather than in the fractions that can be classified as plasmid or virus (Fig.4A-E). It is not possible to tell if the observed significantly fewer >4kb sequences from Falkland Island samples seen in Fig2B and are due to a biological phenomenon or because assembly of equivalent long sequences in FK samples are obscured by a larger diversity of very small sequences present in these samples.

### Genetic content of elements smaller than 2kb

Because very few sequences shorter than 2kb could be assigned to either plasmids or viruses, it is particularly interesting to decipher genetic content and possible biological functions of this biological dark matter. The gene density is similar between the two size fractions <2kb and >2kb, as both have just over 1 complete predicted gene pr kb sequence (1.26genes/kb and 1.18 genes/kb, data not shown). Of these, however there is a remarkable difference as only 29% of genes from small sequences <2kb can be assigned to PFAM whereas this is the case for 45% of genes from >2kb sequences. This difference a likely the result of research focus on larger sequences, leaving genes on smaller sequences unexplored.

PFAMs are averagely encountered 1.99 times in <2kb sequences and almost double that on larger sequences (3.80 encounters/PFAM), indicating an extremely diverse population of genes on the small <2kb sequences. It is known that transposases can create circular intermediate sequences (Sekine, Aihara and Ohtsubo, 1999; Nielsen *et al.*, 2013), and more transposases are found in short sequences than in long ones, despite the <2kb fraction only having 1/3 of the PFAM annotated genes that the longer >2kb fraction have (63 vs 50 genes respectively, data not shown). Further confirming that the small sequences <2kb are not spontaneous single-cell anomalies, the average encounters of <2kb and >2kb sequences are similar, with an average of 1.18 and 1.28 encounters per sequence, meaning that every ca fourth and fifth sequence is encountered more than once.

### Plasmid replication gene diversity

In an effort to find differences between the types of plasmid replication genes present in the two environments, we summarized the fraction of sequences carrying each replication domain (Fig.6A). Samples fluctuate between 8% and 57% of sequences encoding a replication gene (‘Sum’ row), but for no plasmid replication PFAM with more than ten non-redundant sequences was there a significant difference between groups of replication genes (data not shown). For both environments, the dominating plasmid replication types are Rep_1, Rep_2, Rep_3, and RepL. While no difference between environments is seen, the total pool of plasmid replication genes adds significantly to the knowledge on the diversity of replication initiation genes. In Fig.7, the genes from this study with significant similarity to the Rep_2 family that is primarily found in Lactobacillales have been aligned with the PFAM seed database of representative sequences (Punta *et al.*, 2012). The resulting dendrogram demonstrates the large added diversity of the sequences from this study (red and green) and reduces the diversity from the SEED database of representative sequences (black) to merely a clade on the new Rep_2 phylogenetic tree. This pattern has been observed previously and demonstrates the power of mobilomics to discover a diversity of plasmids that have never before been recognized (Jørgensen *et al.*, 2014). This pattern can be seen for all categories of plasmid replication proteins (Appendix 8). Interestingly, Rep_2 genes from Falkland Island (green) and hospital sewer (red) samples do not separate into different clades but are completely shuffled in the tree, a pattern observed for all plasmid replication PFAMs. This point to plasmid diversity being established well before the two populations of rats became separated presumably hundreds of years ago, and subsequently maintenance of plasmid populations.

**Fig. 7.**
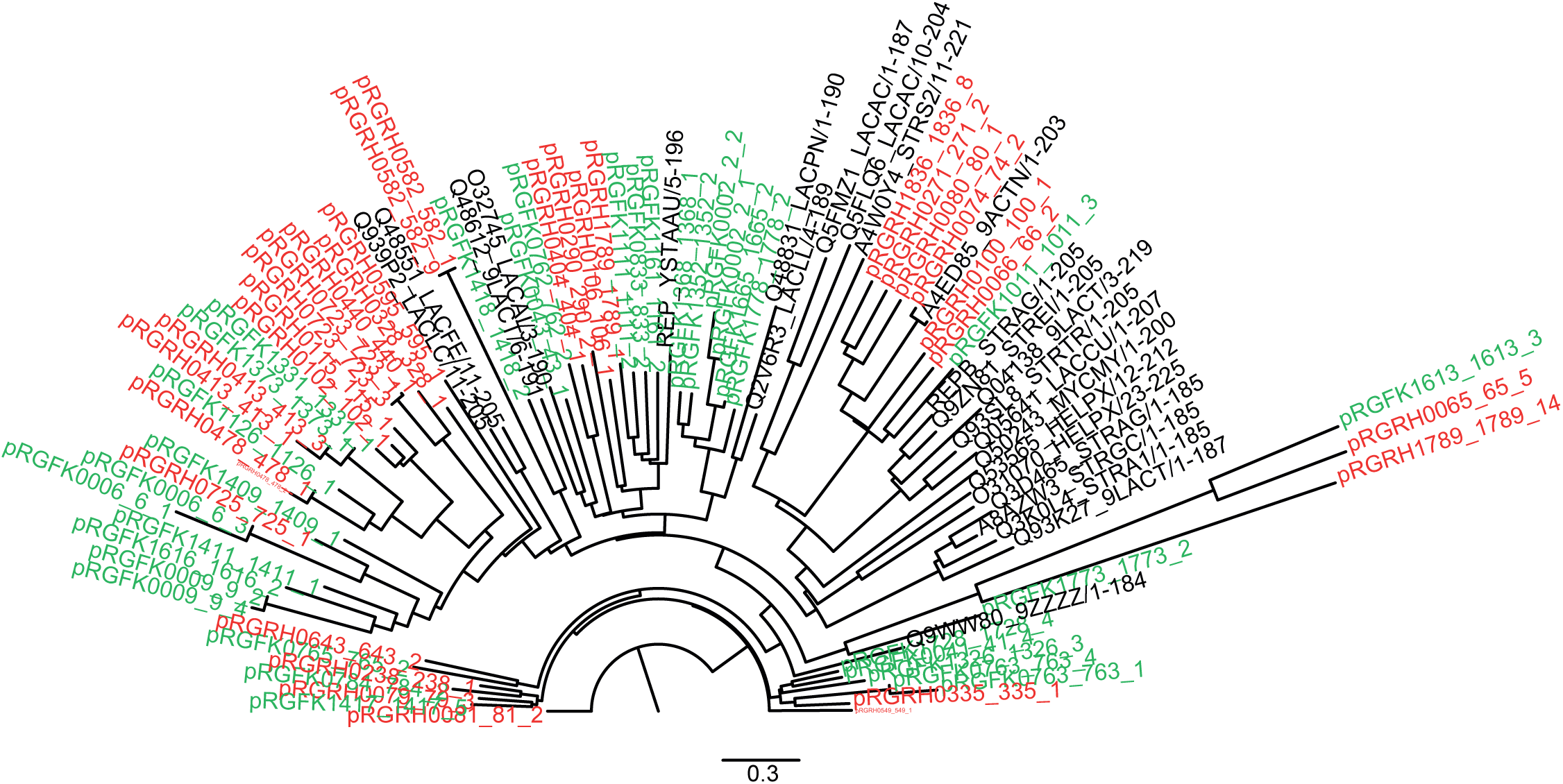
Phylogeny of Rep_2 genes from hospital sewer samples (red), Falkland Island samples (green), and the SEED database of representative sequences on nucleotide level. The neighbor-joining tree is constructed after trimming of sequences to remove sequence length based phylogenic inferences. Notice that entire clades and outgroups exist, reducing the reference sequences to one branch on the dendrogram and illustrating the massive contribution from sequences identified in this study to widely used databases such as the SEED database.

### Diversity of antibiotic resistance genes

Antibiotic resistance is an important aspect of plasmid biology and even though resistance genes are mostly seen on larger plasmids and often in clusters, we wanted to identify antibiotic resistance gene candidates in the dataset of 1869 plasmids. Using a newly published HMM based pipeline (Gibson, Forsberg and Dantas, 2014), we see a larger diversity of ABR genes in hospital sewer sequences, even though they stem from fewer samples (5 vs. 7). We identified 39 potential antibiotic resistance genes of 26 different Resfams families in the 810 hospital sewer sequences and 15 potential antibiotic resistance genes of 11 different Resfams families in the 1,093 Falkland Islands sequences (Table 1). The main class of resistance genes is efflux pumps, accounting for almost half of identified sequences. While it may seem surprising that a substantial amount of ABR genes are found on sequences from the pristine Falkland islands, we speculate that the actual function of the genes may well be different from antibiotic resistance, particularly since efflux pumps are often non-specific excretion systems(Hansen *et al.*, 2007). Because wastes from infectious diseases wards and operating rooms flow through the hospital sewer and hospital sewage has been found to carry substantial amounts of antibiotic compounds (Fredskilde, Nielsen and I/S, 2008), we expected a much higher diversity of ABR genes from these samples, as also seen in (Czekalski *et al.*, 2012). Presumably, the bulk of antibiotic resistance genes are located on plasmids outside of the size range of current mobilomics, why relatively few genes are found. In the search for ABR genes, a shortcoming of assembly of mobilome derived reads should be highlighted: ABR genes are supposed to be rapidly spread through populations when exposed to selective pressure, but a gene existing on two different plasmids will cause them both to break in the assembly process and not be detected.

**Table 1.** Overview of types of antibiotic resistance genes and number of instances detected by Resfams.

| Envir<br>on-<br>ment | total<br>instan<br>ces | Betalacta<br>mases |  |  | efflux<br>pumps |  |  | others |  |  |
| --- | --- | --- | --- | --- | --- | --- | --- | --- | --- | --- |
|  |  | type | resfa<br>m | Instan<br>ces | type | resfa<br>m | instan<br>ces | type | resfa<br>m | instan<br>ces |
| FK | 15 | - | - | 0 | msbA | RF01<br>07 | 7 | Chlor_Acetyltrans_<br>CAT | RF00<br>50 | 8 |
|  |  |  |  |  | macB | RF00<br>89 |  | vanR | RF01<br>54 |  |
|  |  |  |  |  | tolC | RF01<br>47 |  | vanS | RF01<br>55 |  |
|  |  |  |  |  | ramA | RF01<br>14 |  | vanT | RF01<br>56 |  |
|  |  |  |  |  | MFS_efflu<br>x | RF01<br>04 |  | tet_ribosomoal_pro<br>tect | RF01<br>35 |  |
|  |  |  |  |  | norA | RF01<br>09 |  |  |  |  |
| RH | 39 | CepA | RF00<br>47 | 5 | ABC_efflu<br>x | RF00<br>07 | 18 | ANT2 | RF00<br>26 | 16 |
|  |  | ClassB | RF00<br>54 |  | emrE | RF00<br>66 |  | Cfr23_rRNA_meth<br>yltrans | RF00<br>48 |  |
|  |  | ClassC-AmpC | RF00<br>55 |  | macA | RF00<br>88 |  | Cfr23_rRNA_meth<br>yltrans | RF00<br>48 |  |
|  |  | mecR1 | RF00<br>94 |  | macB | RF00<br>89 |  | Chlor_Acetyltrans_<br>CAT | RF00<br>50 |  |
|  |  |  |  |  | MFS_efflu<br>x | RF01<br>04 |  | Erm23S_rRNA_me<br>thyltrans | RF00<br>67 |  |
|  |  |  |  |  | msbA | RF01<br>07 |  | Fluor_Res_DNA_T<br>opo | RF00<br>74 |  |
|  |  |  |  |  | RND_efflu<br>x | RF01<br>15 |  | tet_ribosomoal_pro<br>tect | RF01<br>35 |  |
|  |  |  |  |  | soxR | RF01<br>21 |  | vanC | RF01<br>51 |  |
|  |  |  |  |  | tet_MFS_e<br>fflux | RF01<br>34 |  | vanH | RF01<br>53 |  |
|  |  |  |  |  | tolC | RF01<br>47 |  | vanR | RF01<br>54 |  |
|  |  |  |  |  |  |  |  | vanS | RF01<br>55 |  |
|  |  |  |  |  |  |  |  | AAC6-I | RF00<br>04 |  |

## Conclusions

In this study, we have identified 1869 circular sequences in rat ceca, of which, 504 are plasmid-like, 120 are virus-like, 10 are rat-derived and 1235 are unclassified plasmoids. By a factor of four, this is the largest collection of complete plasmids from a single study to date, and the first and largest collection of complete plasmoids that cannot be classified as plasmid or virus from multiple samples. After previously confirming the validity of the applied methodology, we find a small but interesting overlap in the presence of plasmids, viruses and plasmoids between rat cecum samples from hospital sewers and the pristine Falkland Islands, suggesting that the observed plasmids and plasmoids are not transient but an unrecognized and widespread biological phenomenon. We have demonstrated significantly different size distributions of all classes of elements between environments with in all cases pristine Falkland Island samples having fewer long sequences, pointing to an environmental influence on extrachromosomal element ecology. The elements have been classified to show that very few elements above 2kb of length is not carrying a known plasmid or virus gene while very few sequences shorter than 2 kb carry a known plasmid or virus-related gene. This novel class of plasmoids is enigmatic and poorly fits the known classes of biological units as they seem to not carry complete systems for self-preservation, possibly relying on spontaneous rise or unknown mechanisms for sustained existence. Hopefully, further research will explore this new world of plasmoids to determine possible biological functions, the temporal stability within cells, and the spread of these elements, changing their status from biological dark matter to more meaningful biological categories.

## Methods

Hospital sewer rats were sampled in a collecting room of a large hospital in Copenhagen, Denmark. Rats were euthanized with pellet gun and the ceca were excised directly after euthanization. Rats from the Falkland Islands were caught in snap-traps and sampled within hours after capture. All samples were stored in RNAlater at -80°C until use.

The mobilomes were prepared as previously described (Jørgensen *et al.*, 2014). Libraries were constructed with the Illumina NexteraXT kit and dual indexing primers. The libraries were then sequenced on an Illumina HiSeq2000 (2x100nt) using TruSeq Paired end Dual Index Sequencing Primer Box (Illumina). All bioinformatics work was done in a UNIX environment running Biopieces (Martin Asser Hansen, unpublished, www.biopieces.org). Both ends of reads were trimmed and filtered as described(Jørgensen *et al.*, 2014). Both orphaned and paired end reads were used for assembly. IDBA-UD was used for assembly of reads into contigs and circular sequences were identified using the two-step approach for Illumina sequences validated in (Jørgensen *et al.*, 2014). Mobilome reads were mapped back on doubled circular sequences using bowtie2 (Langmead and Salzberg, 2012) with standard parameters including end-to-end alignment to ensure that short sequences with relatively large end regions was not underrepresented. Comparisons between samples were done with an adapted BLAST script that disregards the observed different breakage points of identical circular elements between samples and only return BLAST hits of full length relative to the query (100nt length difference allowed across the entire sequence, see Appendix xx for script). Known circular sequences was identified using the same script against a downloaded database consisting of the NCBI plasmid collection between 1 and 40 kb (2,940 sequences, downloaded 2015-01-31) and circular sequences in the NCBI nucleotide collection identified with the following search parameters: ‘circular 1000:40000[SLEN]’ yielding 11,458 sequences in the size range 1-40kb (downloaded 2015-01-31). While many of the 11,458 sequences was most definitely not of plasmid, bacterial or even circular origin, this was disregarded since only circular, plasmid-like sequences was found by BLAST searching as described above. Genes were predicted with the Prodigal software and the metagenomics switch (– p meta). Assignment of predicted genes to HMM protein domains were done with Rpsblast (version 2.2.22) and the PFAM database with standard parameters. To remove random hits, a cutoff e-value of 10E-4 was then applied. Chromosomal DNA was identified and quantified as previously described (Jørgensen *et al.*, 2014). Samples with a calculated chromosomal fraction above 20% were excluded from analysis, and most samples had 0-5% chromosomal, 16S rDNA carrying DNA.

Mapping of circular sequences on the rat genome was done by hashing all circular sequences into 100nt bits using the split_seq Biopiece that was then mapped on Rnor6.0 complete *R. norvegicus* genome in CLCgenomics workbench and visual identification of all mapping sites. This approach ensured simultaneously that the breakpoint of the circular sequence did not influence the mapping and disclosed information in repetitive sequences; if a circular sequence had been in a repetitive sequence, the circular sequence bits would have been randomly distributed in repeats and not places shoulder by shoulder as observed for all nine sequences.

PFAM27.0 SEED databases was downloaded from http://pfam.xfam.org/ (February, 2015) for each plasmid replication PFAM and used to construct phylogenetic trees the following way: SEED sequences and sequences form this study was aligned using CLC genomics 7.5.1 with the following parameters: gap open cost: 10, gap extension cost: 1, end gap cost: as any other). The alignment was trimmed to only contain shared regions between samples and realigned. Neighborjoining trees were constructed with standard parameters and 500 bootstraps. Figtree 1.4.2 was used for visualizations.

## Acknowledgements

We thank Karin P. Vestberg and Anette H. Løth for excellent technical assistance and the entire Copenhagen Municipal Pest Control unit for the capture of the hospital sewer specimens. Further, we thank the following institutions for financial support: The Lundbeck Foundation (www.lundbeckfoundation.com), grant project DKnrR44-A4384, the European Union’s 7^th^ Framework Program for Research and Technological Development (grant agreement no. 222625 METAEXPLORE, EUFP7ThemeKBBE-2007-3-3-05) and the Community Research and Development Information Service (CORDIS). The funders had no role in study design, data collection and analysis, decision to publish, or preparation of the manuscript.

## References

1. Lopez-Bueno A, Tamames J, Velazquez D et al. High diversity of the viral community from an Antarctic lake. Science (80-) 2009;326:858–61.

2. Jørgensen TS, Xu Z, Hansen MA et al. Hundreds of Circular Novel Plasmids and DNA Elements Identified in a Rat Cecum Metamobilome. PLoS One 2014;9:e87924.

3. Kav AB, Sasson G, Jami E et al. Insights into the bovine rumen plasmidome. Proc Natl Acad Sci 2012;109:5452–7.

4. Minot S, Bryson A, Chehoud C et al. Rapid evolution of the human gut virome. Proc Natl Acad Sci 2013;110:12450–5.

5. Sachsenröder J, Braun A, Machnowska P et al. Metagenomic identification of novel enteric viruses in urban wild rats and genome characterization of a group A rotavirus. J Gen Virol 2014:vir–0.

6. Barkay T, Smets BF. Horizontal gene flow in microbial communities. ASM NEWS-AMERICAN Soc Microbiol 2005;71:412.

7. Sørensen SJ, Bailey M, Hansen LH et al. Studying plasmid horizontal transfer in situ: a critical review. Nat Rev Microbiol 2005;3:700–10.

8. Gillings MR. Evolutionary consequences of antibiotic use for the resistome, mobilome and microbial pangenome. Front Microbiol 2013;4.

9. Martínez JL, Coque TM, Baquero F. What is a resistance gene? Ranking risk in resistomes. Nat Rev Microbiol 2014.

10. Carattoli A. Plasmids in Gram negatives: molecular typing of resistance plasmids. Int J Med Microbiol 2011;301:654–8.

11. Carattoli A, Bertini A, Villa L et al. Identification of plasmids by PCR-based replicon typing. J Microbiol Methods 2005;63:219–28.

12. Brolund A, Franz+®n O, Melefors +ûjar et al. Plasmidome-analysis of ESBL-producing Escherichia coli using conventional typing and high-throughput sequencing. PLoS One 2013;8:e65793.

13. Jones B V, Marchesi JR. Transposon-aided capture (TRACA) of plasmids resident in the human gut mobile metagenome. Nat Methods 2006;4:55–61.

14. Klümper U, Riber L, Dechesne A et al. Broad host range plasmids can invade an unexpectedly diverse fraction of a soil bacterial community. ISME J 2014.

15. Kav AB, Benhar I, Mizrahi I. A method for purifying high quality and high yield plasmid DNA for metagenomic and deep sequencing approaches. J Microbiol Methods 2013;95:272–9.

16. Walker A. Welcome to the plasmidome. Nat Rev Microbiol 2012;10:379.

17. Sentchilo V, Mayer AP, Guy L et al. Community-wide plasmid gene mobilization and selection. ISME J2013;7:1173–86.

18. Li LL, Norman A, Hansen LH et al. Metamobilomics -expanding our knowledge on the pool of plasmid encoded traits in natural environments using high-throughput sequencing. Clin Microbiol Infect 2012;18:5–7.

19. Kraberger S, Arg++ello-Astorga GR, Greenfield LG et al. Characterisation of a diverse range of circular Rep-encoding DNA viruses recovered from a sewage treatment oxidation pond. Infect Genet Evol 2015.

20. Lanza VF, de Toro M, Garcill+ín-Barcia MP et al. Plasmid flux in Escherichia coli ST131 sublineages, analyzed by plasmid constellation network (PLACNET), a new method for plasmid reconstruction from whole genome sequences. PLoS Genet 2014;10:el004766.

21. Summers D. *The Biology of Plasmids*. Wiley-Blackwell, 1996.

22. Finan TM, Weidner S, Wong K et al. The complete sequence of the 1,683-kb pSymB megaplasmid from the N2-fixing endosymbiont Sinorhizobium meliloti. Proc Natl Acad Sci 2001;98:9889–94.

23. Barnett MJ, Fisher RF, Jones T et al. Nucleotide sequence and predicted functions of the entire Sinorhizobium meliloti pSymA megaplasmid. Proc Natl Acad Sci 2001;98:9883–8.

24. Burian J., Stuchl-¦¦ük S, Kay WW. Replication control of a small cryptic plasmid of Escherichia coli. J Mol Biol1999;294:49–65.

25. Yasukawa H, Hase T, Sakai A et al. Rolling-circle replication of the plasmid pKYM isolated from a gram-negative bacterium. Proc Natl Acad Sci 1991;88:10282–6.

26. Novick RP, Clowes RC, Cohen SN et al. Uniform nomenclature for bacterial plasmids: a proposal. Bacteriol Rev 1976;40:168–89.

27. Rinke C, Schwientek P, Sczyrba A et al. Insights into the phylogeny and coding potential of microbial dark matter. Nature 2013;499:431–7.

28. Dawkins R. The Selfish Gene. Oxford university press, 2006.

29. Jørgensen TS, Kiil AS, Hansen MA et al. Current strategies for mobilome research. Name Front Microbiol 2015;5:750.

30. Krupovic M, Cvirkaite-Krupovic V. Towards a more comprehensive classification of satellite viruses. Nat Rev Microbiol 2012;10:234.

31. Robinson R. Genetics of the Norway rat. 1965.

32. Hilton GM, Cuthbert RJ. Review article: the catastrophic impact of invasive mammalian predators on birds of the UK Overseas Territories: a review and synthesis. Ibis (Lond 1859) 2010;152:443–58.

33. Himsworth CG, Parsons KL, Jardine C et al. Rats, cities, people, and pathogens: a systematic review and narrative synthesis of literature regarding the ecology of rat-associated zoonoses in urban centers. Vector-Borne Zoonotic Dis 2013;13:349–59.

34. Meerburg BG, Singleton GR, Kijlstra A. Rodent-borne diseases and their risks for public health. Crit Rev Microbiol 2009;35:221–70.

35. Tabak MA, Poncet S, Passfield K et al. Invasive species and land bird diversity on remote South Atlantic islands. Biol Invasions 2014;16:341–52.

36. Tabak MA, Poncet S, Passfield K et al. Rat eradication and the resistance and resilience of passerine bird assemblages in the Falkland Islands. J Anim Ecol 2015;84:755–64.

37. Yang MG, Manoharan K, Young AK. Influence and degradation of dietary cellulose in cecum of rats. J Nutr 1969;97:260–4.

38. Montgomery LARR, Macy JM. Characterization of rat cecum cellulolytic bacteria. Appl Environ Microbiol 1982;44:1435–43.

39. Czekalski N, Berthold T, Caucci S et al. Increased levels of multirésistant bacteria and resistance genes after wastewater treatment and their dissemination into Lake Geneva, Switzerland. Front Microbiol 2012;3.

40. Jacobsen L, Wilcks A, Hammer K et al. Horizontal transfer of tet (M) and erm (B) resistance plasmids from food strains of Lactobacillus plantarum to Enterococcus faecalis JH2-2 in the gastrointestinal tract of gnotobiotic rats. FEMS Microbiol Ecol 2007;59:158–66.

41. Norman A, Riber L, Luo W et al. An improved method for including upper size range plasmids in metamobilomes. PLoS One 2014;9:el04405.

42. Li W. Analysis and comparison of very large metagenomes with fast clustering and functional annotation. BMC Bioinformatics 2009;10:359.

43. Storlazzi CT, Lonoce A, Guastadisegni MC et al. Gene amplification as double minutes or homogeneously staining regions in solid tumors: origin and structure. Genome Res 2010;20:1198–206.

44. Altschul SF, Madden TL, Sch+ñffer AA et al. Gapped BLAST and PSI-BLAST: a new generation of protein database search programs. Nucleic Acids Res 1997;25:3389–402.

45. Sekine Y, Aihara K, Ohtsubo E. Linearization and transposition of circular molecules of insertion sequence IS3. J Mol Biol 1999;294:21–34.

46. Nielsen TK, Xu Z, Gözdereliler E et al. Novel Insight into the Genetic Context of the cadAB Genes from a 4-chloro-2-methylphenoxyacetic Acid-Degrading Sphingomonas. PLoS One 2013;8:e83346.

47. Punta M, Coggill PC, Eberhardt RY et al. The Pfam protein families database. Nucleic Acids Res 2012;40:D290–301.

48. Gibson MK, Forsberg KJ, Dantas G. Improved annotation of antibiotic resistance determinants reveals microbial resistomes cluster by ecology. ISME7 2014:1–10.

49. Hansen LH, Jensen LB, S+©rensen HI et al. Substrate specificity of the OqxAB multidrug resistance pump in Escherichia coli and selected enteric bacteria. J Antimicrob Chemother 2007;60:145–7.

50. Fredskilde JWL, Nielsen U, l/S DKEAL. Mâleprogram Pô Rigshospitalet., 2008.

51. Langmead B, Salzberg SL. Fast gapped-read alignment with Bowtie 2. Nat Methods 2012;9:357–9.

